# Genome assembly and association tests identify interacting loci associated with vigor, precocity, and sex in interspecific pistachio rootstocks

**DOI:** 10.1101/2022.06.28.498047

**Authors:** William Palmer, Ewelina Jacygrad, Sagayamary Sagayaradj, Keri Cavanaugh, Rongkui Han, Lien Bertier, Bob Beede, Salih Kafkas, Deborah Golino, John Preece, Richard Michelmore

## Abstract

Understanding the basis of hybrid vigor remains a key question in crop breeding and improvement, especially for rootstock development where F_1_ hybrids are extensively utilized. Full-sibling UCB-1 F_1_ seedling rootstocks are widely planted in commercial pistachio orchards that are generated by crossing two highly heterozygous outbreeding parental trees of *Pistacia atlantica* (female) and *P. integerrima* (male). This results in extensive phenotypic variability, prompting costly removal of low-yielding small trees. To identify the genetic basis of this variability, we assembled chromosome-scale genome assemblies of the parental trees of UCB-1. We genotyped 960 UCB-1 trees in an experimental orchard for which we also collected multi-year phenotypes. We genotyped an additional 1,358 rootstocks in six commercial pistachio orchards and collected single-year tree size data. Genome-wide single marker association tests identified loci associated with tree size and shape, sex, and precocity. In the experimental orchard, we identified multiple trait-associated loci and a strong candidate for ZZ/ZW sex chromosomes. We found significant marker associations unique to different traits and to early vs. late phenotypic measures of the same trait. We detected two loci strongly associated with rootstock size in commercial orchards. Pseudo-testcross classification of markers demonstrated that the trait-associated alleles for each locus were segregating in the gametes of opposite parents. These two loci interact epistatically to generate the bimodal distribution of tree size with undesirable small trees observed by growers. We identified candidate genes within these regions. These findings provide a foundational resource for marker development and genetic selection of vigorous pistachio UCB-1 rootstock.

## Introduction

Pistachio (*Pistacia vera* L., n=15; Anacardiaceae) has been cultivated in the Middle East for thousands of years (Yi et al., 2008); however, the first commercial harvest in the US was not until 1976. Pistachio orchards in the US consist of a clonally propagated scion of *P. vera* grafted onto a non *P. vera* rootstock with 99% of the industry based in California (Kallsen et al., 2009). The most commonly used rootstock, UCB-1, is a F_1_ hybrid of specific *P. atlantica* (female) x *P. integerrima* (male) trees that was established in the 1980s for vigor, resistance to the fungal pathogen Verticillium, and cold tolerance (Ferguson et al., 2005). Because the same two parental trees are always crossed each year to produce UCB-1 seed, all UCB-1 rootstocks are full F_1_ siblings. Over 350,000 acres of UCB-1 trees are planted in California’s Central Valley; these millions of trees represent a unique opportunity for large-scale genetic analysis of phenotypes in hybrid woody species.

Due to the interspecific nature of the cross between outbreeding and highly heterozygous parents, UCB-1 F_1_ seedlings exhibit extensive phenotypic and genetic variability (Jacygrad et al., 2020); the frequency of seedlings exhibiting small size, abnormal branching, and unusual leaf color can be as high as 30% (Ferguson, 2000). The genetic basis of such variability has not been previously investigated. Uneven vigor and stunting are of particular concern to pistachio growers because smaller trees result in decreased nut yield and/or quality (Beede, 2017). As a result, nurseries commonly rogue 10 to 15% of their UCB-1 seedlings based on early growth parameters and visual inspection in an attempt to obtain more vigorous seedlings. However, as we previously reported, growth during the first year is a poor predictor of growth in later years (Jacygrad et al., 2020). Consequently, growers frequently rogue small trees, often several years after planting and at significant cost. This is especially problematic due to the long juvenility period of pistachio, with up to seven years until first flowering and production of nuts (Ferguson, 1998).

The long juvenile period, plus the years necessary to evaluate yield phenotypes, makes conventional pistachio breeding a lengthy process. Consequently, progress in genetic improvement has been slow. Previous genetic work has largely focused on the nut producing scion of *P. vera*. A genetic linkage map for *P. atlantica* has been produced using an F_1_ cross of *P. vera* and *P. atlantica* using a two-way pseudo-testcross strategy, (Turkeli and Kafkas, 2013). High-quality reference genomes are an indispensable tool for accelerating crop genetics. In the Anacardiaceae, published genomes include mango, *Mangifera indica* (Wang et al., 2020) and *P. vera*, although the latter was highly fragmented and not chromosome scale (L50=949.2 kb) (Zeng et al., 2019).

To investigate the genetic basis of tree size and vigor and to provide a foundational resource for selection of pistachio rootstock, we generated chromosome-scale genome assemblies and high-density F_1_ linkage maps of the *P. atlantica* and *P. integerrima* parents of UCB-1. We found strong synteny between each species. We identified one chromosomal scaffold in *P. integerrima* and two in *P. atlantica* with suppressed recombination consistent with the sex-determining Z and W chromosomes (Sola-Campoy et al., 2015). We genotyped 2,358 UCB-1 rootstocks using genotyping-by-sequencing (GBS; Figure 1). Of these, 960 were ungrafted and growing in an experimental orchard at the University of California, Davis; these trees were phenotyped over multiple years and for multiple traits. The remaining 1,398 were grafted and located in six commercial orchards in California. These were phenotyped for both trunk and scion diameter in a single year. Single marker association tests identified multiple trait-associated loci in the experimental orchard. Of these, one locus was detected across multiple years and phenotypes that explained a large proportion of the variance in tree size and correlated traits and is therefore considered to be controlling tree vigor. In the commercial orchard dataset, we only identified two loci, both of which were also detected in the experimental orchard dataset. These loci interact epistatically and explain the bimodal size distribution observed by growers.

**Figure 1.**
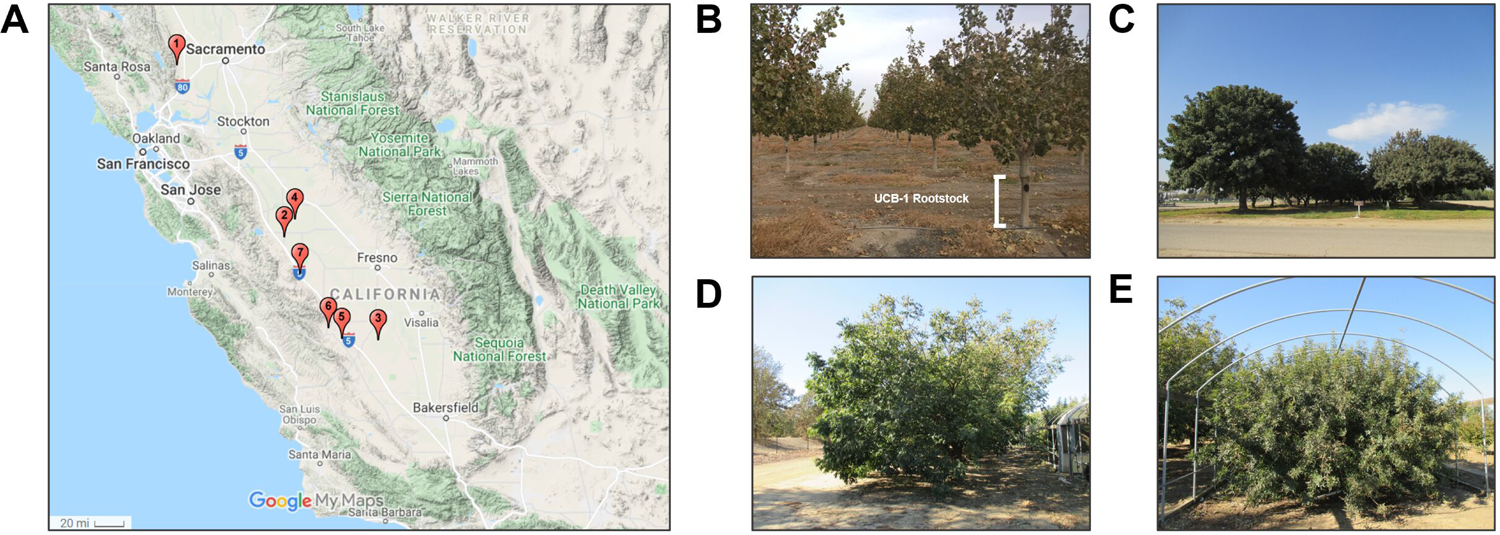
A) Locations of orchards studied in the California Central Valley. In total, 2,358 UCB-1 rootstocks were phenotyped and genotyped. B) Example of a grafted tree in a commercial orchard. The UCB-1 rootstock is highlighted. C) Original trees for *P. integerrima* (left) and *P. atlantica* (right) parental germplasm. D) and E) Clonal parental trees of *P. integerrima* and *P. atlantica* respectively at Foundation Plant Services, Davis.

## Materials and Methods

### Phenotyping of non-grafted and grafted UCB-1 trees

Trees were phenotyped in both an ungrafted experimental orchard at UC Davis Russell Ranch (38°33′00.4″ N, 121°51′26.9″ W) as well as six commercial orchards located in the Central Valley of California (see Table 2). The establishment of the experimental orchard and methods for phenotypic measurements have been described previously (Jacygrad et al. 2020). The sex of 479 UCB-1 trees was determined between February and April 2018–2021 in the experimental orchard at UC Davis Russell Ranch.

### Sample collection, DNA Extraction, and library preparation

#### DNA extraction from F_1_ UCB-1 rootstocks

For plants in the experimental orchard, 0.05 g of young leaves were harvested in the spring, placed in 2 mL Safe-Lock Eppendorf tubes with two 2.3 mm diameter stainless steel grinding beads, flash frozen in liquid nitrogen, and stored at −80°C. For commercial orchards, bark samples (approximately 5 cm^2^) were collected from the rootstock using a hammer and paint scraper. The tools were cleaned with 70% ethanol between the collection of each sample to prevent cross contamination. The samples were kept on ice while transported to the laboratory. The cambium was separated in the lab using a scalpel that was cleaned with 70% ethanol between samples. Cambial samples (0.05–0.1 g) were placed in Eppendorf tubes with stainless steel grinding beads, flash frozen in liquid nitrogen, and stored at −80°C.

Frozen samples were homogenized using a TissuLyser II (Qiagen, Hilden, Germany) set at 16 Hz/min for 2 min. Tools were kept cold during the grinding process. Bark samples that were not successfully homogenized were ground for an additional 2 min or until the tissue turned into a powder. To improve this process, we added one more bead. Homogenized samples were stored at −80°C.

Initially, the Qiagen DNeasy Plant Mini Kit (Qiagen, Hilden, Germany) was used for a small number of samples following the manufacturer’s protocol, with some modifications. Specifically, 450 µL Buffer AP1 was added and 600 µL of the mixture (lysate and Buffer AW1) was transferred into a DNeasy Mini spin column placed in a 2 mL collection tube. Then, we added 30–80 µL of Buffer AE for elution and incubated the sample for 10 min at room temperature (15–25°C). The sample was centrifuged at 20,000 x g (14,000 rpm).

Qiagen DNeasy 96 Plant Kit (Qiagen, Hilden, Germany) was used for the majority of samples following the manufacturer’s protocol with some modifications. Specifically, we followed the protocol in 2 mL Safe-Lock Eppendorf tubes as described in the previous paragraph. After centrifugation, we transferred 400 μL of the supernatant to a collection plate and followed the manufacturer’s protocol for plates. We used collection plates instead of S-Blocks and Elution Microtubes RS racks for all subsequent steps. For elution, we added 30–50 µL Buffer AE and incubated the plates for 10 min at room temperature (15–25°C).

#### DNA and RNA extraction from parental trees (P. integerrima and P. atlantica)

For construction of 10X Genomics and Oxford Nanopore libraries, in the spring and summer, young leaves were harvested in 50 mL Falcon tubes until the entire tube was filled. Then, the collected samples were placed on ice packs while transported to the laboratory. Before placing samples in a −80°C freezer, they were flash frozen with liquid nitrogen.

For construction of Dovetail Hi-C libraries, in the spring and summer, young leaves (3–4g) were harvested in 50 mL Falcon tubes until the entire tube was filled, and then the samples were shipped overnight in an insulated container with dry ice to Dovetail Genomics.

#### Strand-Specific RNA Libraries

For construction of RNA seq libraries, fully developed leaves were harvested in 50 mL Falcon tubes and placed in liquid nitrogen while transported to the laboratory, where they were transferred to a −80°C freezer.

All leaves collected in 50 mL Falcon tubes (except those sent to Dovetail Genomics) were ground in liquid nitrogen using a pre-chilled mortar and pestle until the frozen leaves were a fine powder.

### Standard DNA extraction (for GBS and Skim-Seq sequencing)

A Qiagen DNeasy Plant Mini Kit (Qiagen, Hilden, Germany) was used for a small number of samples following the manufacturer’s protocol, with some modifications. Specifically, 450 µL Buffer AP1 was added and 600 µL of the mixture (lysate and Buffer AW1) was transferred into a DNeasy Mini spin column placed in a 2 mL collection tube. Then, we added 30–80 µL of Buffer AE for elution and incubated the sample for 10 min at room temperature (15–25°C). The sample was centrifuged at 20,000 x g (14,000 rpm).

Qiagen DNeasy 96 Plant Kit (Qiagen, Hilden, Germany) was used for the majority of samples following the manufacturer’s protocol, with some modifications. Specifically, we followed the protocol in 2 mL Safe-Lock Eppendorf tubes as described in the previous paragraph. After centrifugation, we transferred 400 μL of the supernatant to a collection plate and followed the manufacturer’s protocol for plates. We used collection plates instead of S-Blocks and Elution Microtubes RS racks for all subsequent steps. For elution, we added 30–50 µL Buffer AE and incubated the plates for 10 min at room temperature (15–25°C).

### High molecular weight DNA extraction

High molecular weight (HWM) extraction for 10X and Oxford Nanopore sequencing was done following the CTAB protocol (Webb and Knapp, 1990), with 1% polyvinyl pyrrolidone (PVP) 40,000 and 1% sodium metabisulfite added to CTAB buffer. CTAB extraction was followed by a Salt:Chloroform Wash (Pacific Biosciences, Menlo Park, California, USA) to clean up the high molecular weight genomic DNA prior to performing the library protocol. The Salt:Chloroform Wash consists of a high salt, low ethanol wash to remove polysaccharides before DNA is precipitated from the solution. The extracted DNA was kept at 4°C overnight. Then, the DNA was quantified using a Qubit. The DNA quality was checked on 0.5% agarose gel. Pulsed field gel electrophoresis was run by the DNA Technologies Core at the UC Davis Genome Center to determine the quantity and quality of the high-molecular-weight DNA.

### RNA extraction

Total RNA, for Illumina sequencing and transcriptome assembly, was extracted with the Qiagen RNeasy® Plant Mini Kit (Qiagen, Hilden, Germany) with RLT buffer optimized for pistachio tissue (∼0.3 g) by adding 1% sodium metabisulfite and 2.5% PVP as a reducing agent instead of ß-mercaptoethanol. Additionally, 210 µL of 20% lauroyl sarkosyl was added to the RLT buffer containing ground tissue, then the sample was heated at 70°C for 10 min before the solution was passed through a QIAshredder. Total RNA was eluted twice from the spin column with RNAse-free water. A Qubit was used to quantify total RNA, and 2% agarose gel was used to assess RNA quality. One hundred micrograms of total RNA was used for mRNA capture. A Dynabeads™ mRNA Purification Kit (for mRNA purification from total RNA preps) was used to isolate mRNA (Thermo Fisher Scientific, Waltham, Massachusetts, USA).

### Library construction and sequencing

### 10X Genomics Libraries

10X Genomics libraries were prepared using 10X Genomics library preparation kit (Pleasanton, CA, US) by the DNA Technologies Core at the UC Davis Genome Center. Libraries were sequenced on an Illumina Hiseq 4000 as PE150.

### Oxford Nanopore Technology Libraries

Oxford Nanopore libraries for the same *P. atlantica* and *P. integerrima* individuals above were prepared using an Oxford Nanopore Technologies library preparation kit (Oxford, United Kingdom) by the DNA Technologies Core at the UC Davis Genome Center. The libraries were sequenced on a PromethION (Oxford Nanopore).

### Strand-Specific RNA Libraries

The Cold Spring Harbor Protocol (Zhong et al., 2011) was used to construct strand specific RNA seq libraries for each parental species using a pool of clonal trees sharing the same genotype as those above. mRNA (400 ng) was used for cDNA synthesis. cDNA was sheared using a Covaris E220 sonicator after completion of the second strand reaction instead of shearing mRNA as in the protocol. PCR was done using KAPA HiFi HotStart ReadyMix PCR Kit (Kapa Biosystems, Wilmington, Massachusetts, US). A microcapillary nucleic acid fragment analysis for quality control of the cDNA library was conducted using a Bioanalyzer, and a Qubit was used for library quantification. Libraries were sequenced on an Illumina Hiseq 2500 as PE150 and Illumina Miseq as PE250. The libraries were sequenced by the DNA Technologies Core at the UC Davis Genome Center. The concentration of pooled barcoded libraries was measured using Qubit; quality control check and quantitation was done using an Agilent Bioanalyzer.

### F_1_ WGS Skim-Seq libraries

Paired-end libraries for *P. atlantica* and *P. integerrima* were prepared using NEBNext End Repair Module (Ipswich, MA, USA), Enzymatics Klenow (3′→ 5′ exo-) (Beverly, MA, USA), Enzymatics T4 DNA Ligase (Rapid) (Beverly, MA, USA), and KAPA HiFi Polymerase MM or Phusion High-Fidelity PCR MasterMix (Kapa Biosystems, Wilmington, Massachusetts, US), Ampure XP beads (Beckman Coulter, Miami, FL, US), and reagents following the manufacturers’ protocols. The protocol generates libraries with an average size of 300–350 bp. DNA was sheared using a Covaris E220 sonicator. Libraries were sequenced on an Illumina Hiseq 3000 as PE150. The concentration of pooled barcoded libraries was measured using a Qubit; quality control check and quantification was done using an Agilent Bioanalyzer. The libraries were sequenced by the DNA Technologies Core at the UC Davis Genome Center.

Illumina (short-read) Whole Genome Shotgun (WGS) libraries were made for 95 ungrafted UCB-1 trees by the DNA Technologies Core at the UC Davis Genome Center using an Illumina TruSeq DNA HT Sample Prep Kit. A Covaris focused sonicator was used to physically fragment the DNA to a narrow size range suitable for Illumina library preparations, and a total of 1 µg of DNA was provided per sample. All the samples were normalized to ±20% of the average sample concentration. DNA was sheared using a Covaris E220 sonicator. Libraries were sequenced on an Illumina Hiseq 3000 as PE150. The concentration of pooled barcoded libraries was measured using a Qubit; an Agilent Bioanalyzer was used for the quality check and quantification. The libraries were sequenced by the IGM Genomics Center at the University of California, San Diego.

### F_1_ GBS Libraries

The libraries were constructed according to Elshire et al. (2011). We used the methylation sensitive AvaII enzyme that was suitable for pistachio.

From leaves, GBS libraries were made for 960 ungrafted UCB-1 trees. Restriction enzyme digestion was conducted using 5.5 µL of 0.6 ng/µL AVAII enzyme and 100 ng DNA at 37°C for 1 hour. Ligation was followed by cleanup conducted with the Omega E.Z.N.A. Cycle Pure Kit (Norcross, GA, USA). The pooled samples were amplified via 16 PCR cycles followed by clean up with the Omega E.Z.N.A. Cycle Pure Kit.

From bark, GBS libraries were made for 1,416 grafted UCB-1 trees. A low input DNA protocol was used for bark samples. Restriction enzyme digestion was conducted using 1.1 µL of 0.6 ng/µL AVAII enzyme and 20 ng DNA at 37°C for 16 hours. Five times the volume of the libraries was pooled and cleaned up with an Omega E.Z.N.A. Cycle Pure Kit.

The concentration of pooled barcoded libraries was measured using a Qubit, and the quality control was checked using an Agilent Bioanalyzer. The libraries were sequenced by the DNA Technologies Core at the UC Davis Genome Center on a single HiSeq 4000 lane with single end 90 bp reads.

### Genome size estimation

Genome size was initially estimated experimentally by Fluorescence-Activated Cell Sorting (FACs). Approximately 0.5 cm^2^ of fresh tissue from young leaves of *Orzya sativa*, a *P. atlantica* female, and a *P. integerrima* male were used. *O. sativa* was used as a reference because it is expected to have a comparable genome size to pistachio (430 Mb).

Sysmex CyStain PI Absolute P Kit reagent kit for nuclei extraction and DNA staining of nuclear DNA from plant tissues was used to determine the absolute and relative genome size and ploidy level of a *P. atlantica* female and *P. integerrima* male individual. The reagent was created specifically for selected configurations of the Sysmex CyFlow Ploidy Analyser and CyFlow Space flow cytometers. Briefly, the samples were co-chopped in 500 μL Nucleic Extraction Buffer. A razor blade with a single edge was used for chopping each sample in a Petri dish. A 50-μm mesh filter was used to filter the resulting homogenate and staining solution was added, and samples were incubated in a dark −20 freezer. CellQuest™ software was used to acquire and analyze data from the flow cytometer.

A computational estimate of genome size and heterozygosity was also calculated. Using Illumina short reads, kmer counts were generated for each parental tree using Jellyfish (Marcais and Kingsford., 2011) and further analyzed with Genomescope (Vurture et al., 2017).

### Assembly

10X reads were initially assembled with Supernova 1.1.4. One *pseudohap2* fasta record was selected and provided to Dovetail for further scaffolding with Dovetail Hi-C reads and HiRise (Putnam et al., 2016). Ultra-long Oxford Nanopore reads (*P. atlantica*, ∼75 Gb, L50 32 kb; *P. integerrima* ∼50 Gb, L50 40 kb) were preprocessed with Porechop (https://github.com/rrwick/Porechop) and Nanofilt with the parameters *L10000, q10, HTcrop50*. Reads were subsequently assembled with SHASTA 0.1.0 (Shafin et al., 2020) and default parameters. ONT reads were mapped back to each assembly with Minimap2 (Li, 2018) and the parameters -ax map-ont. These alignments were used for error correction with Margin Polish (https://github.com/UCSC-nanopore-cgl/MarginPolish). Short WGS Illumina reads for each species were additionally mapped to each assembly using bwa-mem, with these alignments used for further error correction with Pilon 1.23 (https://github.com/broadinstitute/pilon). As the resulting contigs fell short of chromosome scale, they were scaffolded and oriented against the above chromosome-scale 10X+Hi-C assemblies using RaGOO v1.1 (Alonge et al 2019). This resulted in large chromosome-scale scaffolds for each species, and a small number of unassigned contigs, which we designated unanchored.

### Assembly validation and correction

Dovetail Hi-C reads for each species were aligned to each assembly using Bowtie2. Hi-C contact matrices were generated with JuicerTools makeTagDirectory and JuicerTools tagDir2hicFile.pl and then visualized with Juicebox 1.11.08 (https://github.com/aidenlab/juicer). In this way, a single, large inter-chromosomal misassembly was identified on Chr 15 of *P. atlantica* (Figure S1). Gap boundaries flanking the feature were identified and the entire sequence was replaced with a gap of 100 bp. Sequence masked in this way was assigned to new contigs and designated unanchored. No evidence of large scale misassembly was observed in *P. integerrima*.

Assemblies were aligned to each other with Nucmer (Mummer4) and delta filter with the options −l 100 -i 95, and alignments visualized with Mummerplot (Kurtz et al. 2004). Based on the alignment of each species to a genome assembly of *P. vera* (Kafkas et al., unpublished) entire chromosomal scaffolds were reverse complemented using revcomp and named to match the existing chromosomal nomenclature for *P. vera*. In the case of *P. atlantica*, two chromosomal scaffolds showed a 2:1 relationship to a single chromosomal scaffold (Chromosome 14) in *P. integerrima* and *P. vera*, which we designated Chromosome 14.1 and 14.2.

Kmer spectra were calculated with KAT Comp and short Illumina reads vs. each assembly. These were visualized with KAT plot spectra-cn (Mapleson et al., 2017). Sequence completeness was assessed with BUSCO V3 and the Embryophyta odb10 database.

### High Density Linkage Mapping and Assembly Correction

A high density linkage map for each species was constructed with LepMap3 (Rastas 2017). Illumina short reads from each parental species were aligned to each assembly with bwa mem and genotypes called with freebayes and the parameters --use-best-n-alleles 2 --strict-vcf --standard-filters. Low coverage (skim-seq) Illumina WGS reads for 94 F_1_ individuals at the UC Davis Experimental Orchard were aligned to each assembly with bwa-mem and genotypes called with freebayes and the parameters --use-best-n-alleles 4 --strict-vcf --standard-filters. A custom script was used to identify and retain only sites in testcross configuration (for each species/map, respectively), remove sites with greater than 10% missing data, segregation distortion outside of chi-square expectation at p=0.05, and finally generate the requisite input table for LepMap3. The map was reconstructed using default parameters, except for the following tools: Filtering2 (data tolerance=0.001); SeparateChromosomes2 (lodLimit=1); JoinSingles2All (lodLimit=3 iterate=1).

The resulting maps identified a small number of contigs misoriented (reverse complemented) relative to the rest of the chromosomal scaffolds in which they were anchored. In these cases, contig boundaries were manually identified and contigs reverse complemented with revcomp and the original sequence (of identical length) replaced.

### Transcriptome Assembly and Gene Annotation with Maker

Prior to gene prediction, each assembly was masked using RepeatMasker v4.0.9 with a custom repeat library built from each assembly using RepeatModeler v2.0 (Flynn et al., 2020).

To annotate the final assembly of each species, we utilized the MAKER-P pipeline (Cantarel et al., 2008) to combine evidence from *ab initio* gene finding, homology-based gene prediction, and two transcriptomes that we generated *de novo*.

*Ab initio* gene finding was performed with Augustus v3.0.3. For homology-based prediction, we downloaded and utilized cDNA and peptide sequences of 15 closely related plant species (see Table S1) from Ensembl plants (https://plants.ensembl.org/index.html). Furthermore a *de novo* transcriptome was generated for each species using RNA-seq reads and Trinity 2.9.1 (Haas et al., 2013) and utilized as input.

### Association testing and statistical analyses

#### GBS Genotyping and Association Testing

GBS reads were preprocessed and demultiplexed with GBSx (Herten et al., 2015). Reads from these 2,358 F_1_ rootstocks were aligned to each assembly with bwa-mem and genotypes called with freebayes and the parameters --use-best-n-alleles 4 --strict- vcf --standard-filters --skip-coverage 1000000 --min-alternate-count 10. VCFtools was used to generate a genotype matrix, remove indels, and remove sites with non-biallelic calls. This genotype matrix was read into R and filtered to remove sites with >100 individuals, as well as combined with a matrix of metadata and phenotypic measures. Variants were classified by the observed segregation pattern of each parent, and variants not in testcross configuration removed. Genotypes were recoded from character (0/0, 0/1) to numeric vectors by replacing a given genotype value (e.g., 0/1) with a numerical assignment in the matrix (e.g., 1). For the sex association testing model, the phenotypes male or female were recoded to 0 or 1 in the same manner, with genotypes again recoded as 0, 1, or 2. Single marker association tests were performed at each single nucleotide polymorphism (SNP) position by fitting a linear model of the form lm(trait value ∼ genotype), whereby phenotype is a vector of trait values, and genotype is the numeric encoding of the F_1_ genotypes. In the commercial orchard dataset, orchard was included as a covariate: lm(trait value ∼ genotype + orchard), where orchard was a factor of orchard assignment. Linear models were implemented in R version 4 with the function lm().

#### Interaction between loci

All statistical tests were performed in R version 4. Kruskal Wallace tests were performed with the function kruskal.test, Wilcox Rank Sum tests with the wilcox.test function, T-tests using t.test(), and ANOVA as implemented in anova() and lm(). Estimates of η^2^ were calculated using the R package LSR (https://github.com/djnavarro/lsr).

#### Identification of Candidate Genes

Regions within 1 Mb of the markers most significantly associated with the trunk diameter phenotype on Chromosomes 3 and 9 were extracted as the QTL intervals for the vigor trait. Predicted gene models within each QTL interval were extracted. Variant calls from each species vs. each assembly were annotated with a predicted function using snpEff (Cingolani et al., 2012). Gene models were annotated with their best hit to Arabidopsis using blastp, the maker peptide prediction for a given gene, and Arabidopsis TAIR (Table S1).

## Results

### Generation of two high-quality reference genome sequences

We generated two high-quality chromosome-scale genome assemblies, one for each of the two parents of UCB-1 rootstock: a *P. atlantica* individual (female) and a *P. integerrima* individual (male), using the precise genetic individuals that are crossed to produce UCB-1 (Figure 2). A combination of Illumina short reads (10X and Hi-C libraries) and Oxford Nanopore long reads were utilized, followed by manual correction against both Hi-C contact matrices and a high-density linkage map. The same workflow was applied to each species separately.

**Figure 2.**
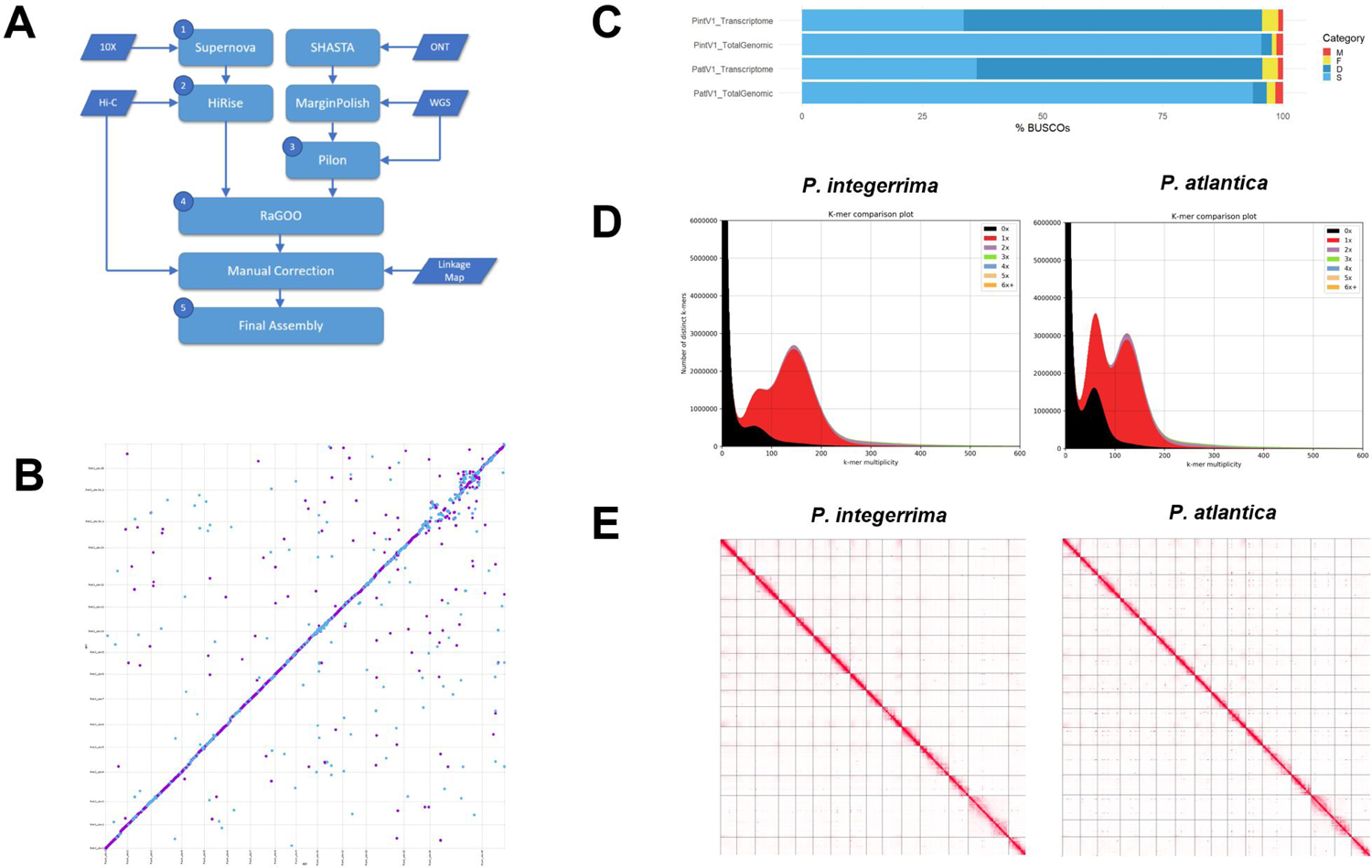
Genome assembly workflow and quality assessment. A) Hybrid assembly workflow incorporating short reads, long reads, and manual correction used to assemble both parental species to chromosome scale. Data sources are presented as parallelograms and processed data as rectangles. Full assembly metrics for each step are presented in Table S3. B) Mummerplot of synteny between assemblies *of P. atlantica* and *P. integerrima*. Weak synteny was only observed on Chromosome 14. C) BUSCO completeness of each assembly and transcriptome. D) KAT kmer spectrum plots for each assembly against short Illumina reads. E) Genome-wide Hi-C contact matrices for both final assemblies: PintV1 and PatlV1.

### Genome size estimation

Previous cytological studies of *P. atlantica* and *P. integerrima* have indicated 14 autosomes and a pair of sex chromosomes (2n=30) (Al-Saghir, 2010). Our initial estimation of genome size and heterozygosity was performed using Genomescope (Vurture et al., 2017) and short read Illumina data. Kmer spectrum analysis (Table S2) predicted a haploid genome size of ∼345 Mb for *P. integerrima* and ∼370 Mb for *P. atlantica*. This is likely an underestimate due to collapsed repeat content. Both repeat length and heterozygosity were estimated to be greater in *P. atlantica* than *P. integerrima*, with an estimated heterozygosity rate of 1.29% vs. 0.64%, and a repeat length of 134 Mb vs. 112 Mb, respectively (Table S2). The heterozygosity rate of *P. atlantica* is comparable to that of mango (1.29% vs. 1.5%; Wang et al., 2020). Genome size was also estimated by FACs using *Arabidopsis* and rice for calibration (Bennett et al., 2003). FACs estimated larger genome sizes of 507 Mb in *P. integerrima* and 560 Mb in *P. atlantica* (Table S2; Motalebipour et al., 2016); these values are more in keeping with the published estimates for *P. vera* (Motalebipour et al., 2016; Zeng et al., 2019) and other *Pistacia* species (https://cvalues.science.kew.org/search/angiosperm).

### Initial short read assembly with 10X Supernova and Dovetail Hi-C HiRise

Initial assemblies were generated using >100 Gb of 10X-Chromium reads assembled with Supernova 1.1.4 (Table S3, step 1). For *P. integerrima*, this yielded 32,291 scaffolds (L50 of 400 kb) and a scaffold length of 459 Mb. For *P. atlantica*, this yielded 52,644 scaffolds (L50 of 190 kb) and a scaffold length of 510 Mb. Gap percent in these scaffolds was 7.7% and 8.2% in *P. integerrima* and *P. atlantica*, respectively (Step 1; Figure 2A, Table S3.1). As the genomic coverage was similar and the assembly approach identical between species, the observed differences in metrics are expected to be related to the differing genome size, repeat content, and heterozygosity of each species.

To improve the contiguity of these initial assemblies, they were further scaffolded using Dovetail Hi-C + HiRise (Putnam et al., 2016). This improved the scaffold L50 of *P. atlantica* to 26 Mb and *P. integerrima* to 24 Mb (Step 2; Figure 2A, Table S3.1). This resulted in 15 large (>10 Mb) super-scaffolds for *P. integerrima* and 16 for *P. atlantica*. This is consistent with the haploid chromosome count expected for a male (14 autosomes + a ZZ pair) and a female (14 autosomes + a ZW pair). However, large gaps remained in each assembly (8.5% for *P. atlantica* and 7.8% for *P. integerrima*). This step was also expected to eliminate contamination, as non-*Pistacia* contigs will not exhibit contact to the primary chromosomal super-scaffolds and so will be removed.

### Assembly with Oxford Nanopore long reads

To improve contiguity and eliminate gaps, independent assemblies were generated for each species using approximately 50 Gb of sequence in *P. integerrima* and 75 Gb for *P. atlantica* from two flow cells of ONT Promethion for each species. These assemblies were error corrected using the raw ONT data followed by Illumina paired-end data. This yielded 1,255 scaffolds (L_50_ of 2 Mb) and a scaffold length of 476 Mb for *P. integerrima*, and 2,694 scaffolds (L_50_ of 700 kb) and a scaffold length of 510 Mb for *P. atlantica* (Step 3; Figure 2A, Table S3.1). As with the 10X Supernova assemblies, the greater scaffold count and lower scaffold L50 observed in *P. atlantica* is likely due to the genome being larger, more repetitive, and more heterozygous than *P. integerrima*, a common challenge of plant genome assembly.

### Combining assemblies using reference guided scaffolding

As the SHASTA scaffold L50 fell short of the very large (>10 Mb) scaffolds attained using 10X + Dovetail Hi-C, we further scaffolded these assemblies using reference guided scaffolding and the 10X+Hi-C assemblies as a reference. This yielded near final assemblies of 33 scaffolds (L50 of 28 Mb) and a length of 475 Mb for *P. integerrima*, and 91 scaffolds (L50 of 30 Mb) and a length of 509 Mb for *P. atlantica*, with a gap percent of 0.029% and 0.05%, respectively (Step 4; Figure 2A; Table S3.1). Sequences not placed on the large chromosomal super-scaffolds were split to contigs and designated unanchored. Unassigned sequence accounted for only 500 kb in *P. integerrima* and 6.7 Mb in *P. atlantica*, or 0.1% and 1.3% of each species’ assembled sequence, respectively. In total, assembled contigs covered 93% of the *P. integerrima* genome size and 90% for *P. atlantica*, as predicted by FACs. A complete list of assembly statistics for each step is available in Table S3.1.

### Hi-C contact matrices and assembly correction

Beyond assembly statistics, the quality and accuracy of each assembly was assessed using multiple approaches. Firstly, a Hi-C contact matrix for each species was generated using Homer and short Illumina reads from the Dovetail Hi-C library. Visualization with Juicer identified a ∼3 Mb region of Chromosome 15 in *P. atlantica* exhibiting poor linkage to neighboring sequence and strong contact with other chromosomal scaffolds (Figure S1). This was considered to be a misassembly, and the three contigs found in this region were removed and demoted to unanchored status. Final construction of the Hi-C contact map following correction supported 15 chromosomal super-scaffolds in *P. integerrima* and 16 in *P. atlantica* with a strong signal of linearity along the length of each (Figure 2E).

### Synteny between P. atlantica and P. integerrima assemblies

To examine synteny, we aligned our *P. atlantica* and *P. integerrima* assemblies against each other. At the 10 kb scale, both assemblies showed an extremely high level of collinearity across all chromosomes, except for Chromosome 14 (Figure 2B). In this case, *P. integerrima* Chr14 showed synteny to two separate chromosomal super-scaffolds in *P. atlantica*, which we denoted Chr14.1 and 14.2. Synteny between Chr14.1/2 and Chr14 is poor, a feature that may be due to the accumulation of differences between species on these non-recombining chromosomes (see Figure 3C, 3D, and Linkage Mapping below) or some degree of chimeric misassembly on chromosomes. To ensure consistency with established nomenclature, chromosomes were named based on alignment and synteny to *P. vera* (S. Kafkas et al., unpublished).

**Figure 3.**
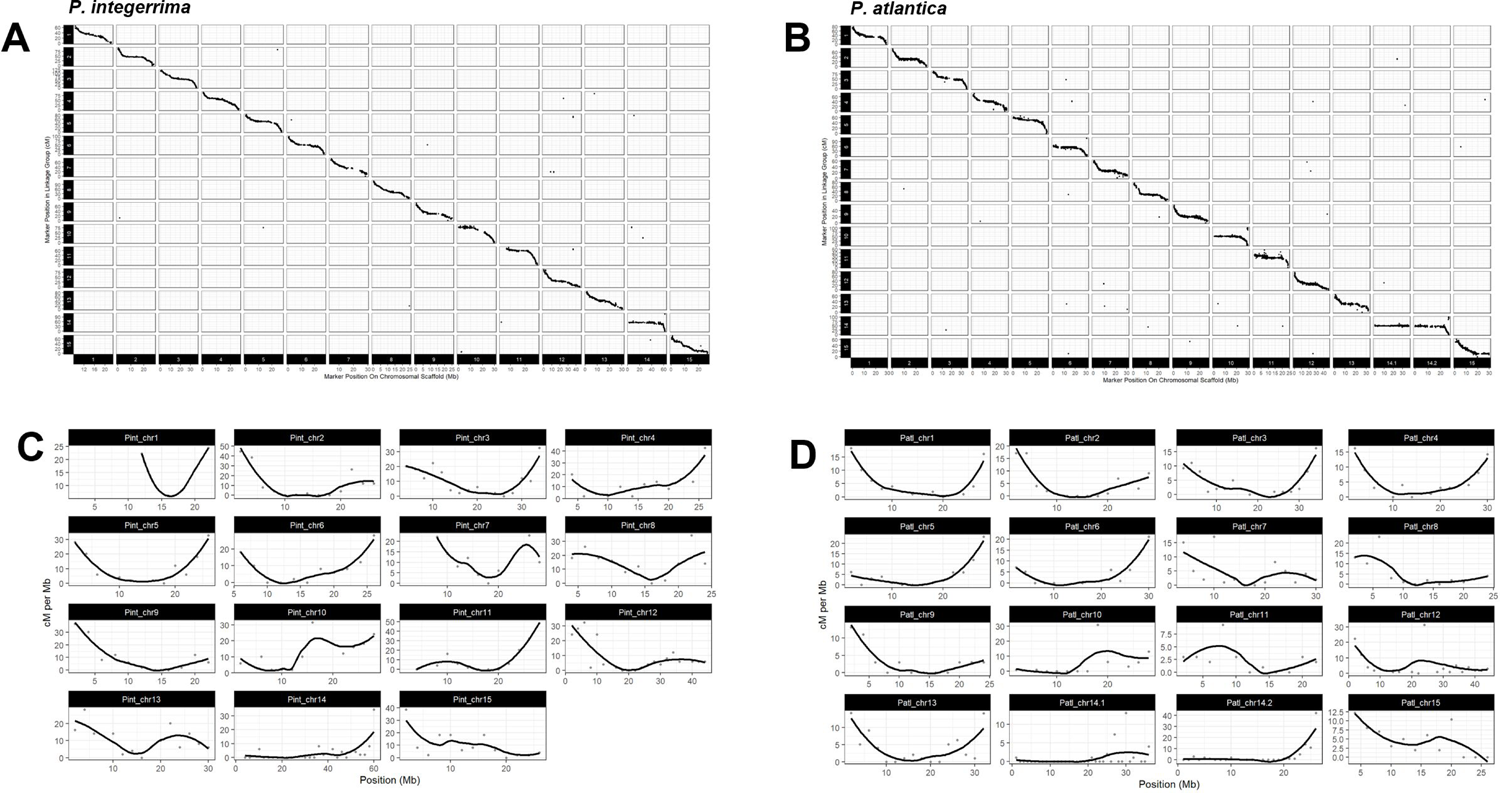
A and B) Marey Maps (physical vs. genetic position) for the position of markers in each assembly vs. the associated linkage map for *P. integerrima* and *P. atlantica*, respectively. C and D) Plots of recombination rate vs. physical position for each chromosome of *P. integerrima* and *P. atlantica*, respectively.

### Assembly Quality Assessment with BUSCO and KAT

The extent to which gene space was reconstructed was examined using BUSCO v2 (Simao et al., 2015). Of 1,375 conserved plant (embryophyta odb10) genes, 97.7% were found to be complete in *P. integerrima* and 96.7% in *P. atlantica*. A slightly lower fraction of genes were both complete and single copy in each case (95.5% and 93.7%, respectively). Complete and duplicated BUSCOs were low at 2.2% in *P. integerrima* and 2.9% in *P. atlantica*. Complete but fragmented BUSCOs totaled 0.9% and 1.7%, with missing BUSCOs at 1.4% and 1.6% in each species (Figure 2C, Table S3.1.2). These results compare favorably to other highly contiguous chromosome-scale plant genomes (summarized in Marks et al., 2021).

Assembly quality was further confirmed using KAT (Mapleson et al., 2015) to test for the presence of kmers in a deep coverage Illumina read set vs. the assemblies. Assembly spectra copy number plots demonstrated the expected distribution of kmer-counts and multiplicity for each assembly, with two peaks representing homozygous and heterozygous content, respectively. The mode of the second peak was ∼2x the multiplicity of the first, and the missing content peak was approximately half the area of the first peak (Figure 2D).

### Transcriptome Assembly and Gene Annotation

To permit more accurate gene model prediction, we initially generated *de novo* transcriptome assemblies for each species using strand-specific RNA-seq data from fully developed leaves. in a transcriptome of 309,845 transcripts in *P. integerrima* and 211,300 in *P. atlantica*; transcript sequences totaled 303 Mb and 236 Mb in each species, respectively (Table S3.1).

Annotation with MAKER resulted in 37,678 predicted gene models in *P. atlantica* and 36,748 gene models for *P. integerrima*, with a total gene length of 93 Mb and 94 Mb, respectively. This corresponded to 18.6% of the genome covered by genes and 6.8% of the genome covered by coding sequence in *P. atlantica*. Similarly in *P. integerrima*, 19.8% of the genome was covered by genes and 7.3% by coding sequence. These 37,678 and 36,748 high quality gene models from *P. atlantica* and *P. integerrima* were further annotated using the best blastp hit to a database of <u>Arabidopsis TAIR 10.1 proteins</u>.

### High density linkage mapping and assembly correction

A high-density F_1_ pseudo-testcross linkage map was constructed using LepMap3 for each parental species using 94 F_1_ UCB-1 ungrafted individuals randomly selected from the experimental orchard at UC Davis sequenced with low-coverage, whole genome Illumina paired-end reads. Maps were constructed separately for each parent by using only SNPs in test-cross configuration and recombining in the species for which the map was being reconstructed (i.e., aaxab where a is the reference allele). Linkage groups (LGs) were numbered (1–15) by their 1:1 correspondence to chromosomal super-scaffolds. LepMap3 was chosen for map reconstruction for its ability to explicitly handle F_1_ testcross data as well as strong performance with a very large (>10,000) set of input markers.

For *P. atlantica*, 35,858 input SNPs were provided to LepMap3, of which 98% (35,239) were assigned to 25 LGs. Of these LGs, LGs 1–15 contained 99.9% (35,203) of the assigned markers and exhibited high log likelihoods ranging from −109806.2296 (LG 1) to −39835.1871 (LG 15). *P. atlantica* chromosomal super-scaffolds 14.1 and 14.2 collapsed to the same linkage group (LG 14) in the map. LGs 16–23 exhibited low log likelihoods ranging from −362.462 to −6.9 and are likely artifactual.

For *P. integerrima*, 12,529 input SNPs were provided to LepMap3, of which 97.5% (12,216) were assigned to 21 LGs. Of these LGs, LGs 1–15 contained 99.7% (12,181) of the assigned markers and exhibited high log likelihoods ranging from −45845.5044 (LG 1) to −8462.3177 (LG 15). As with *P. atlantica,* LGs 16–21 exhibited low log likelihoods ranging from −540.2631 to −6.9078 and are likely artifacts.

Plotting linkage map marker order vs. physical position in the genome assembly demonstrated strong collinearity of each genome assembly with the genetic data (Figure 3A, 3B). However, several contigs were identified in this approach that appeared to be misoriented relative to the rest of their respective chromosomal scaffolds. Such contigs were split at gap positions (“N”), manually reverse complemented, and re-inserted at the same position. Furthermore, a small number of isolated individual markers exhibited interchromosomal linkage grouping. It is likely these represent repetitive sequences or fine scale misassemblies. No large-scale interchromosomal groupings of markers were observed.

Recombination rate in *P. integerrima* was higher than in *P. atlantica* (1,334/507 and 1,204/560 cM/Mb respectively). To examine the recombination rate within each species, each chromosome was divided into 1 Mb bins with centiMorgans counted across each. In both *P. atlantica* and *P. integerrima*, most chromosomal super-scaffolds exhibited elevated recombination towards their distal ends, with decreased recombination around the center, creating a characteristic parabolic or u-shaped curve (Figure 3C, 3D). This is expected and likely due to suppressed recombination around the centromere of each chromosome. In *P. atlantica*, Chromosome 14.1 had a rate consistently at or near zero along its entire length. A similar pattern was observed on Chromosome 14.2; however, a recombining region of ∼10 Mb was observed towards one end. In *P. integerrima*, Chromosome 14 also exhibited little to no recombination. These results are consistent with Chromosomes 14.1 and 14.2 in *P. atlantica*, and Chromosome 14 in *P. integerrima* being the sex chromosomes, which are expected to be a non-recombining pair (Sola-Campoy et al., 2015). Complete marey map data may be found in supplementary tables S4.1 and S4.2.

### Phenotyping of ungrafted (experimental) and grafted commercial F_1_ UCB-1

In total, we phenotyped 2,358 F_1_ UCB-1 rootstocks using manual approaches. This adds additional years of data to the phenotyping dataset of Jacygrad et al. (2020) for the ungrafted experimental orchard, as well as data for two further commercial orchards containing both younger and older rootstocks than those reported in the previous phenotyping study.

In the ungrafted experimental orchard at UC Davis, 960 trees were phenotyped over seven years, as previously described in Jacygrad et al. (2020). After year three, every other tree was removed to prevent overlap, reducing the count in subsequent years to 479. For each tree, four morphometric traits were recorded: trunk diameter, tree height, canopy height, and canopy width (Figure 4A). Due to minimal canopy of young seedlings, canopy traits were not measured in the first year. Trait measurements in the experimental orchard were generally found to be normally or skew-normally distributed and both the mean and variance of each was found to increase with age (Figure 4A). At least some collinearity was found to exist between most pairs of traits (Figure 4B), with correlation between measures in later years generally positive and strong (R^2^>0.7). Trait values in earlier years were found to be a poor predictor of trait values in later years (R^2^<0.5), as previously reported (Jacygrad et al., 2020).

**Figure 4.**
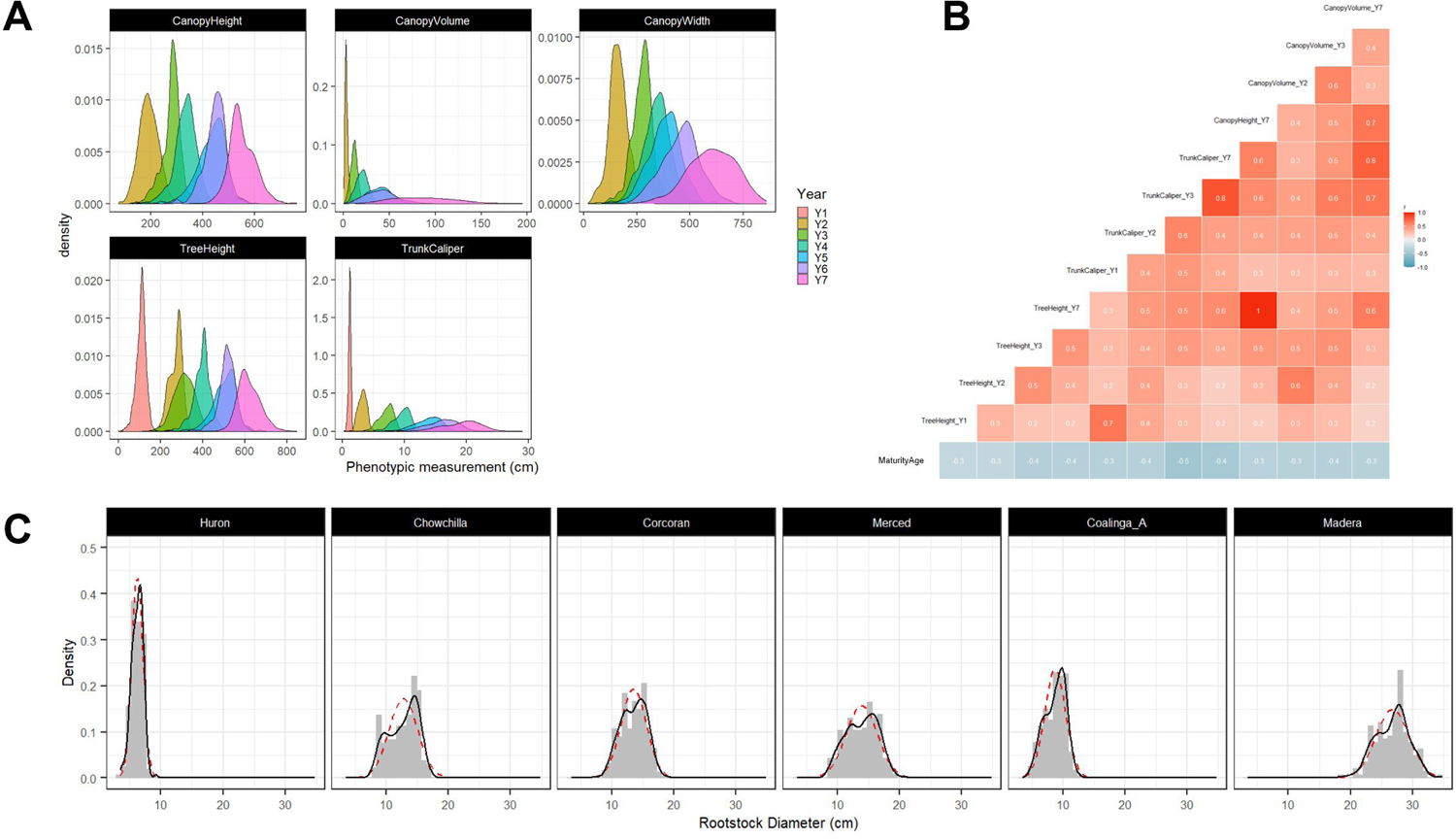
Phenotyping of 960 trees in an ungrafted experimental orchard and 1,358 trees across seven commercial pistachio orchards. A) Phenotypic distribution of five traits across multiple years in the ungrafted experimental orchard. B) Correlation matrix for the 13 experimental orchard traits used for subsequent association t-testing. C) Rootstock trunk diameter size distributions in the six commercial orchards studied. Dotted red line indicates an idealized normal distribution with the same mean and standard deviation; black line indicates a kernel density estimate of the data. Panels are organized left to right by orchard age (youngest to oldest).

The age of maturity or precocity (defined as years until first flowering) and sex were recorded as trees matured. In the experimental ungrafted orchard, precocity was found to be weakly to moderately negatively correlated with all traits (R^2^=-0.3 to −0.5), consistent with more vigorous trees attaining larger size faster and flowering earlier (Figure 4B). As *Pistacia* trees do not typically flower until years 5 to 8, tree sex was only recorded for the 440 remaining trees that had flowered by year 8 (2021), identifying 240 males and 200 females. The sex of ungrafted trees in the experimental orchard was determined by manual examination of floral morphology. Trees started flowering in 2018, with 93.3% of trees having flowered by 2021 (year 8). There was a small yet significant difference in precocity between male and female trees with the mean years to maturity being 6.48 for males and 6.61 for females (t=-2.468, p=0.014).

In the six commercial orchards, trunk diameter for the UCB-1 rootstock and *P. vera* scion were measured as previously described (Jacygrad et al., 2020). Rootstock and scion diameter were strongly correlated (R^2^=0.93, Figure S2). Consistent with our previous study (Jacygrad et al., 2020), a strong bimodality of trunk diameter was observed in all but one orchard (Figure 4C). This bimodality of size is consistent with reports by growers of stunted trees. Unlike the other five orchards studied, where two distinct modes of size were evident, trunk diameter was observed to have low variance and be skew-normally distributed in the three-year-old orchard at Huron (Figure 4C, first panel). This is most readily explained by the trees at Huron being juvenile and the smallest and youngest trees in the dataset (Table 2).

For the association testing described below, we reduced our final phenotyping dataset to those with a correlation coefficient (R^2^) below 90% because correlations between annually collected traits are expected to produce redundant results in association testing. Consequently, in the experimental orchard dataset, we retained measures for only years 1, 2, 3, and 7. For canopy traits, we retained only canopy volume and not its individual components of width and height. In the commercial orchard dataset, trunk but not scion diameter was retained (Figure 4). In total, this resulted in 12 individual traits in the experimental orchard dataset and one in the commercial orchard dataset (Table S2). The complete phenotyping dataset can be found in Table S5.

### Genotyping of UCB-1 rootstock

All F_1_ UCB-1 rootstocks phenotyped in the experimental and commercial orchards were genotyped using genotyping-by-sequencing (GBS). In total, 6.8 billion 100 bp Illumina single-end GBS reads were generated across 2,358 F_1_ UCB-1 individuals. Alignment and SNP/genotype calling against each reference identified 10,292 variant positions using the *P. atlantica* reference sequence and 10,445 variant positions using the *P. integerrima* reference. Only sites in testcross configuration and segregating approximately 1:1 (chi-square deviation from 1:1 at p=0.05) were retained. This reduced the final dataset to 3,229 F_1_ SNPs called against *P. integerrima* and 2,924 called against *P. atlantica* for use in association testing of the experimental and commercial orchards.

### Association mapping of the sex chromosomes

To identify the sex associated chromosomes of UCB-1 rootstock, we performed genome-wide single marker association testing using our GBS-derived genotypes and sex assignment based on floral morphology of 440 F_1_ individuals at the UC Davis experimental orchard. *Pistacia* spp. are dioecious and sex is thought to be determined by a ZW system, whereby males are homogametic (carrying two Z chromosomes) and females are heterogametic (carrying one Z and one W chromosome) (Kafkas et al., 2015; Sola-Campoy et al., 2015). Therefore, males (ZZ) may exhibit less polymorphism on the sex chromosome compared to females in the F_1_ (ZW). Tests were performed utilizing genotypes called by aligning F_1_ reads against each reference genome separately and data were combined in single marker association (SMA) tests at each SNP position using a fixed effects regression model. This identified two chromosomal scaffolds in the female *P. atlantica* assembly and one homologous chromosomal scaffold in the male *P. integerrima* assembly strongly associated with sex. Of the markers identified using the *P. atlantica* reference, 48 SNPs were significantly associated with sex (p=0.05 corrected for multiple testing by Bonferroni correction)(Figure 5B). Of these, 45 out of 48 (93%) were associated with either Chromosome 14.1 or 14.2. Both scaffolds exhibited little recombination in the linkage map. These data are consistent with 14.1 or 14.2 of the female assembly, representing much of the Z and W haplotypes or these chromosomal scaffolds being assembly chimeras. Similar results were obtained using markers identified using the *P. integerrima* reference; 49 SNPs were significantly associated with sex (p=0.05 Bonferroni corrected). Of these, 41 out of 49 (84%) were found only on chromosomal superscaffold 14 (Figure 5A). The small number of hits on other chromosomes likely represent shared repeat content and/or fine scale misassembly. The significant hits on Chromosome 14 were distributed across a large region (∼20 Mb to ∼38 Mb), were highly significant (-log10(p)=9 to 187, Bonferroni significance threshold=5.7), and had large R^2^ values (R^2^_max_=0.85) (Figure 5A). Chromosome 14 exhibited little recombination along most of its length and therefore, any sex determining locus is likely to be in strong linkage with the rest of the chromosome (Figure 3C, 3D). A complete list of significant hits can be found in Table S6.1 and S6.2.

**Figure 5.**
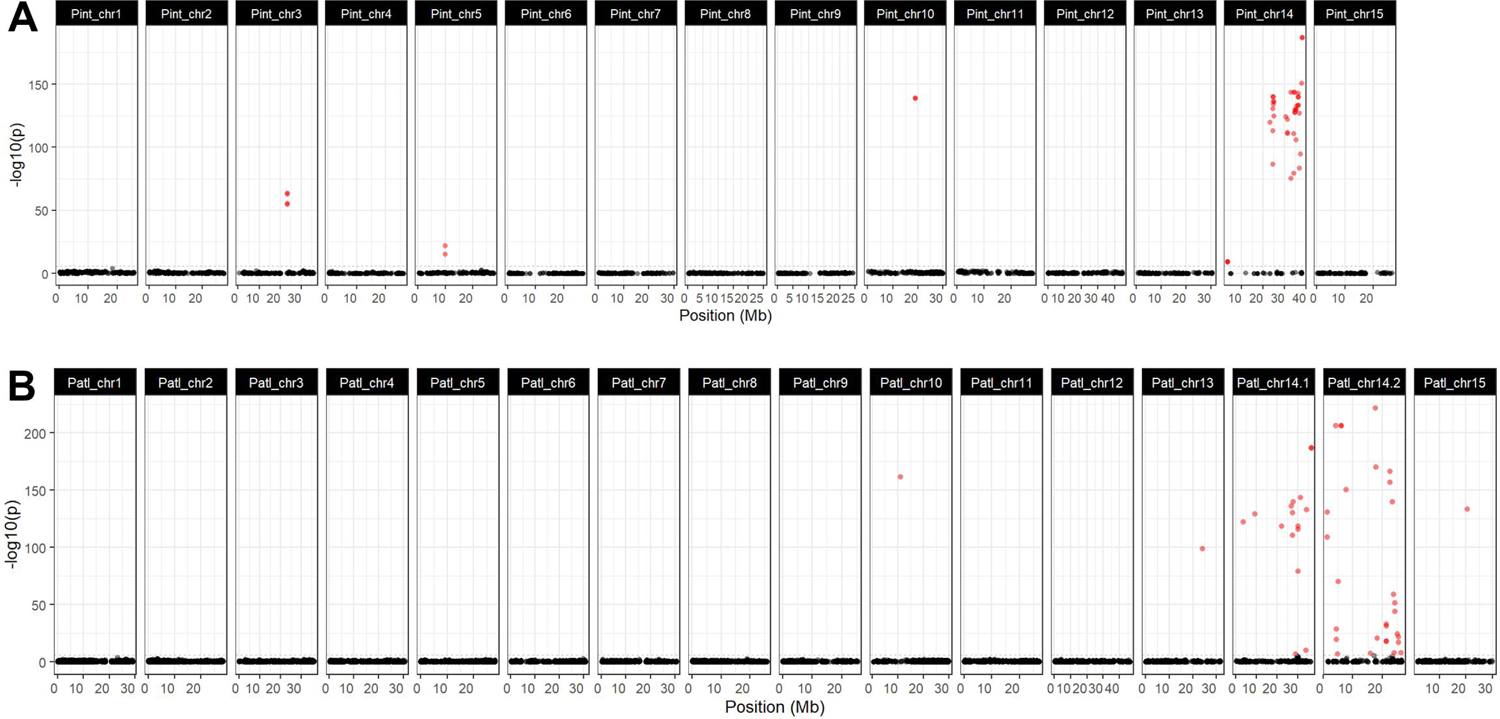
Manhattan plots of association tests between sex and genotype. Results obtained with each of the two genome assemblies independently are shown. A) *P. integerrima*; B) *P. atlantica*. Significant hits (p<0.05 Bonferroni corrected) are shown in red; non-significant in black.

### Genome-wide association tests of multi-year phenotyping in the ungrafted experimental orchard

To identify loci associated with several traits in the experimental orchard dataset, we performed genome-wide single marker association testing by applying a fixed effect regression model to the F_1_ GBS-derived genotypes and phenotypic values for each trait collected. Due to the experimental orchard having a much greater sample size, being ungrafted, and measured multiple times over seven years, we analyzed the experimental orchard and commercial orchards separately, fitting a different model in each case (Table 1). In the case of the experimental orchard with multi-year sampling, each year was treated and examined as an independent trait. In total, this resulted in 13 independent genome-wide association tests across four traits: precocity (a single measure of years to maturity), tree height and trunk diameter in four separate years (years 1, 2, 3, and 7), and canopy volume in three separate years (years 2, 3, and 7). All tests were performed independently against each reference genome.

**Table 1.** Details of models and sample sizes applied in association testing.

| Orchard | Tree Count | Phenotypes | Model |
| --- | --- | --- | --- |
| Experimental ungrafted UCB-1 | 960 (years 1–3)<br>478 (year 7)<br>440 (precocity, by year 8) | 13 total: trunk diameter, tree height, canopy volume (years 1–3, 7).<br>Precocity recorded from years 5 to 8. | lm(trait value ~ genotype) |
| Experimental ungrafted UCB-1 | 240 Male<br>200 Female | Sex (male vs. female) | lm(trait value ~ genotype) |
| 6x Commercial, grafted UCB-1 (see Table 2) | 1398 | Trunk diameter | lm(trait value ~ genotype + orchard) |

Of the SNPs called against the *P. integerrima* assembly, 242 were significantly associated with at least one of the 13 phenotypes (p<0.05 Bonferroni corrected; Figure 6A). Significant associations were detected on 11 out of 15 chromosomes. Peaks of significant SNPs on Chromosomes 8, 9, 11, 13, and 15 were well defined and contained clusters of highly significant markers. A small region centered around 200–600 kb on Chromosome 9 contained highly significant associations with all traits except tree height in year 1. Pint_chr9:245436 had the largest -log10p value reaching -log10p=36.87 for trunk diameter in year 3; the same SNP also had the largest R^2^ values with the greatest being 22% for trunk diameter in year 7 (-log10p=26.99). On Chromosome 3, two distinct clusters of significant markers were observed at opposite ends of the chromosome. The first region, centered around ∼6 Mb, had the most significantly associated markers for trunk diameter and canopy volume across multiple years. Only markers for tree height and trunk diameter in year 1 had maximal significance in the second region towards the end of Chromosome 3. Although highly significant, the R^2^ for markers in either peak ranged from only 2% to 6%. Precocity (age at first flowering) was associated with only two regions, the previously described locus at the start of Chromosome 9 and a second region on Chromosome 8. Some markers were only significant for year 1 data; clusters of highly significant markers for trunk diameter in year 1 were associated with Chromosomes 8 and 15 (Figure 6A). The finding that a unique and different set of loci are associated with year 1 phenotypes fits the finding that traits measured in the first year are poorly correlated with measures in later years (Jacygrad et al., 2020).

**Figure 6.**
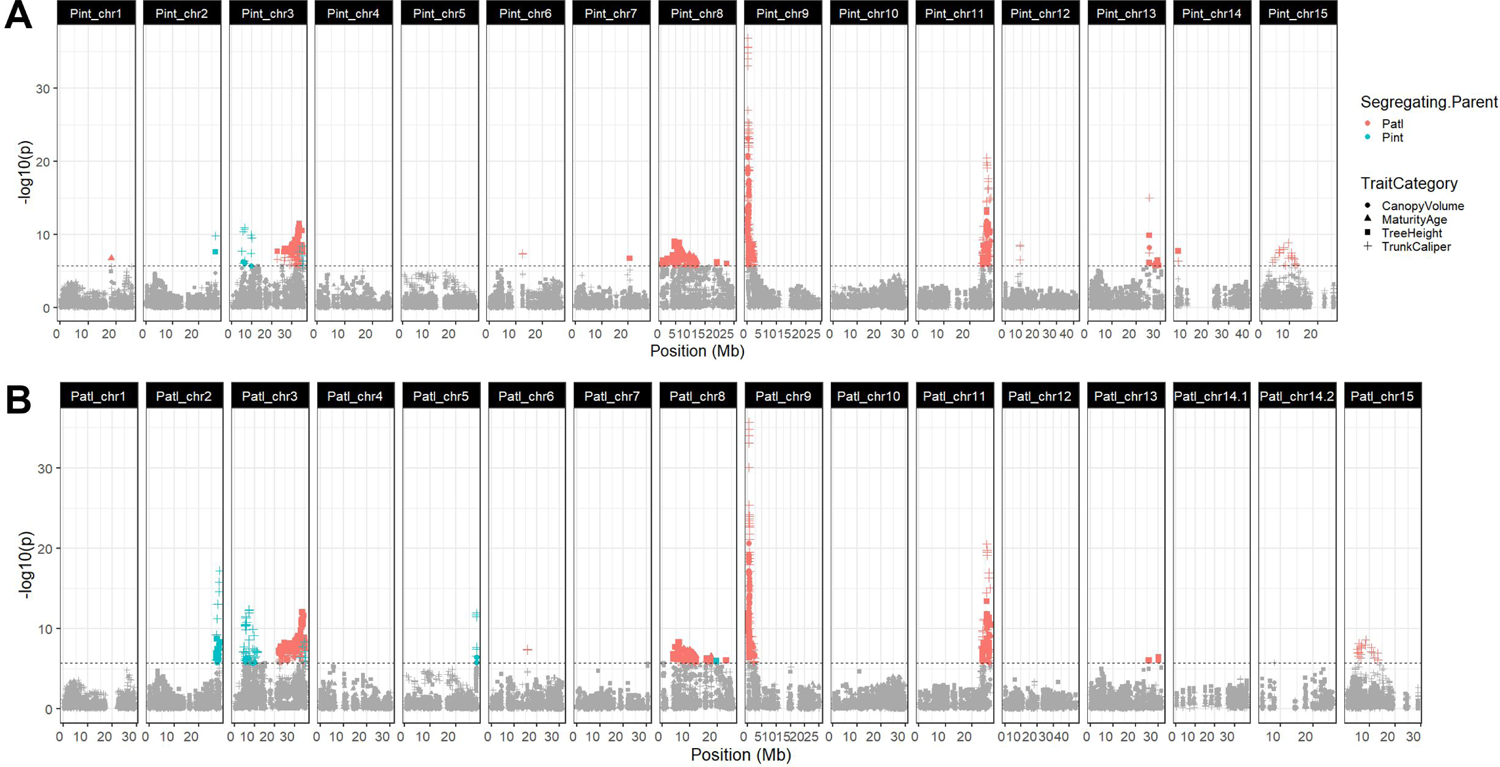
Combined Manhattan plots of trait associations for 13 traits in four categories (canopy volume: circle; precocity: triangle; trunk diameter: cross; tree height: square). Significant hits, as determined by Bonferroni correction, are colored by segregating parent (*P. atlantica*: red; *P. integerrima*: blue). Results obtained with SNPs called against each of the two genome assemblies independently are shown. A) *P. integerrima*. B) *P. atlantica*.

Of the SNPs called against the *P. atlantica* assembly, highly similar but non-identical results were observed, with 312 SNPs significantly associated with at least one of the 13 phenotypes (p<0.05 Bonferroni corrected; Figure 6B). Significant associations were detected on nine out of the 16 chromosomes. As with the *P. integerrima* reference, clear peaks of associated markers were found on Chromosomes 8, 9, 11, and 15 and observed to be in syntenic locations. On *P. atlantica* Chromosomes 2 and 5, a large number of trunk diameter associated markers were observed compared to only a small number on *P. integerrima* Chromosome 2, and none on *P. integerrima* Chromosome 5. As with the *P. integerrima* reference, two distinct groupings of markers were observed on Chromosome 3, with the second again associated with trunk diameter and tree height in year 1, as well as canopy volume in year 2. All markers on Chromosome 15 were again only associated with trunk diameter in year 1.

While many markers were highly significant, their R^2^ values were typically low regardless of the trait (median R^2^ <0.04). Only markers associated with trunk diameter in years 2, 3, and 7, and canopy volume in year 7 had an R^2^ greater than 10% at any given SNP and all of these were located within the small region at the start of Chromosome 9 (Table S7).

We were able to identify in which parental set of gametes each SNP was segregating because we genotyped each parent. Most clusters of markers were exclusively heterozygous in *P. atlantica*; this is consistent with the *P. atlantica* parent having much greater heterozygosity (see *Genome Size Estimation* above and Table S2). Only markers with associations on Chromosome 2, 3, 5, and 8 were heterozygous in *P. integerrima* (Table S7). On Chromosome 3, markers associated with trunk diameter (years 2, 3, and 7) and canopy volume (years 3 and 7) were segregating in the gametes of *P. integerrima* and located in the first 10 Mb of the chromosome. A second locus containing markers exclusively associated with trunk caliper in year 1 was also segregating in the gametes of *P. integerrima* and located in the last 10 Mb of the chromosome. A small number of markers segregating in the gametes of *P. integerrima* located towards the end of Chromosome 3 were associated with canopy volume in year 2 only.

### Genome-wide association tests of single-year phenotyping in six grafted commercial orchards

To identify loci associated with trunk diameter in six commercial orchards (Tables 1 and 2), we performed genome-wide single marker association testing by applying a fixed effect regression model to the F_1_ GBS-derived genotypes and trunk diameter measures for 1,398 trees. To control for environmental variation as well as the different ages/scion cultivars between orchards, we also included “orchard” as a covariate in the model. Tests were performed using SNPs called against each reference genome independently.

**Table 2.** Experimental and commercial orchards sampled

| Figure 1A | Orchard location | Coordinates (lat, long) | Status | Age of trees (years) | Scion cultivar | Number of trees |
| --- | --- | --- | --- | --- | --- | --- |
| 1 | Davis | 38.550107, -121.857605 | Experimental Orchard (Ungrafted). | Multi-year morphometric sampling from 1 year old to 7 years old. Precocity measured until year | Ungrafted | 960 (Years 1–3)<br>470 (Years 4–7) |
|  |  |  |  | 8. |  | 440<br>(Precocity) |
| 2 | Chowchilla | 37.128744, -<br>120.575375 | Commercial orchard<br>(Grafted with *P. vera* scion) | 7 | *P. vera* cv. Golden Hills | 164 |
| 3 | Corcoran | 36.097520,-<br>119.598992 |  | 8 | *P. vera* cv. Golden Hills | 291 |
| 4 | Merced | 37.156753,-<br>120.556466 |  | 8 | *P. vera* cv. Kerman | 287 |
| 5 | Coalinga | 36.1015374,-<br>120.0066809 |  | 8 | *P. vera* cv. Golden Hills | 290 |
| 6 | Huron | 36.196442, -<br>120.153580 |  | 3 | *P. vera* cv. Golden Hills | 192 |
| 7 | Madera | 36.682587, -<br>120.4787 |  | 18 | *P. vera* cv. | 192 |
|  |  | 97 |  |  | Kerman |  |

Of the SNPs called against the *P. integerrima* reference, 74 variants were found to be significantly associated with trunk diameter (p<0.05 Bonferroni corrected; Figure 7A), whereas 68 were significantly associated when using SNPs called against the *P. atlantica* reference (p<0.05 Bonferroni corrected; Figure 7B). All 74 SNPs called against *P. integerrima* and 62 of the 68 SNPs called against *P. atlantica* were significantly associated within the first 1 Mb of Chromosome 9 and the first 10 Mb of Chromosome 3. These were the same highly significant associations detected with data from the experimental orchard. In addition, using the SNPs called against the *P. atlantica* assembly, a small number of significant markers were identified on Chromosomes 11 and 13 (Figure 7).

**Figure 7.**
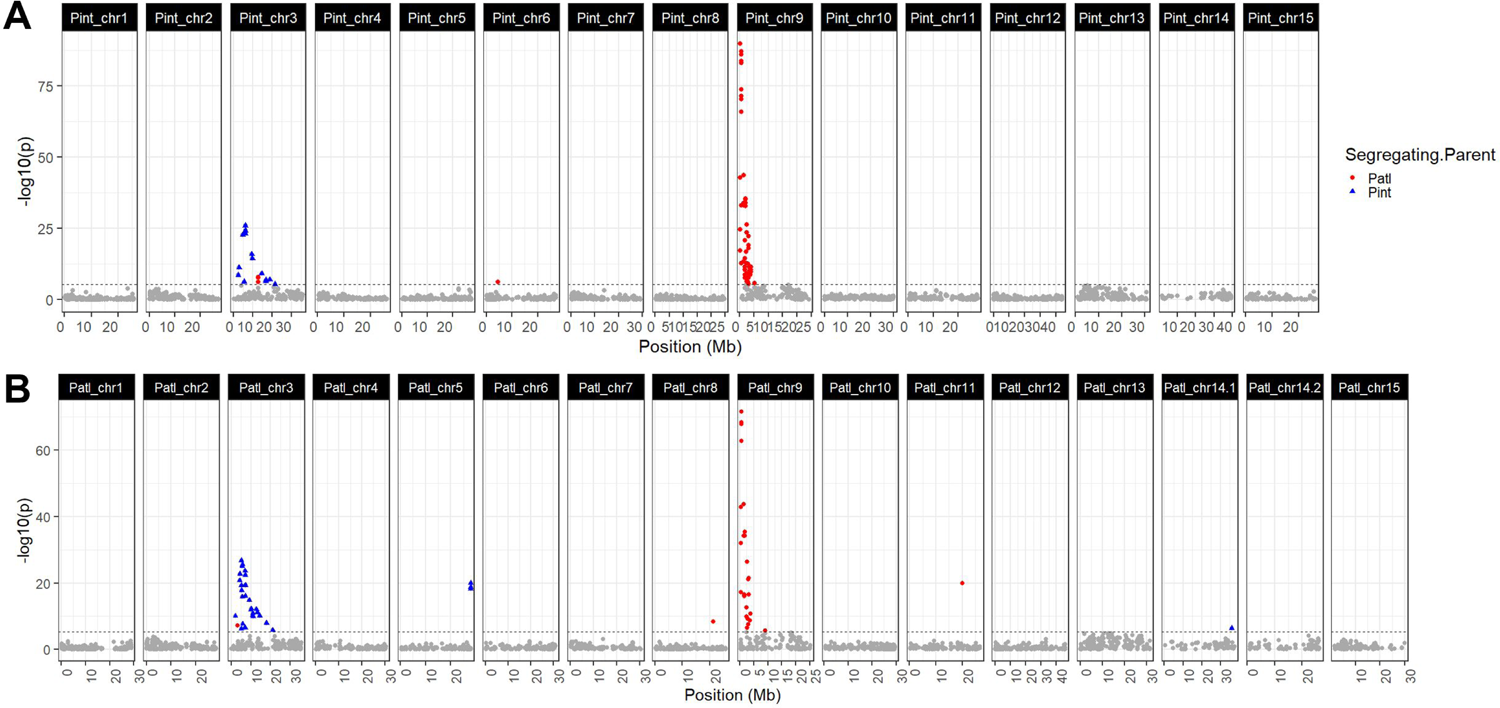
Manhattan plots of single marker association tests for trunk diameter. Significant hits, as determined by Bonferroni correction, are colored by segregating parent (*P. atlantica*: red; *P. integerrima*: blue). Results obtained with SNPs called against each of the two genome assemblies are shown. A) *P. integerrima*. B) *P. atlantica*.

The significance of markers on Chromosome 9 was much greater than that of markers on Chromosome 3 (Table S8). In addition, the proportion of variance explained by the most significant marker on each chromosome (R^2^ or η^2^) was greater on Chromosome 9 for most orchards, with Corcoran notably deviating from this trend (Table S9.2). Furthermore, the proportion of variance in trunk diameter explained by the genotype varied between orchards. For example, the most significant SNP called against *P. atlantica* on Chromosome 9 was found to explain 45% of the variance in trunk diameter at Chowchilla but only 4.4% at Corcoran (Table S9.2). The most significant SNP on Chromosome 9 called against *P. integerrima* explained 51% of the variance at Chowchilla but only 6.4% at Corcoran. Results for Chromosome 3 similarly varied, with the proportion of variance explained by the most significant SNP observed to be 14% and 15% at Chowchilla (*P. atlantica, P. integerrima* called SNPs, respectively) and 5.4% and 6.5% at Corcoran (Table S9.2).

The significant peaks for trunk diameter on Chromosomes 3 and 9 were concordant with those identified in the experimental orchard in years 3 and older. As with data from the experimental orchard, markers on Chromosome 3 segregated in the gametes of *P. integerrima* and markers on Chromosome 9 in the gametes of *P. atlantica.* Clear peaks associated with trunk diameter in the experimental orchard on Chromosomes 2, 5, 11, and 15 were not observed with the commercial orchard data (Figure 6 and 7; Tables S7, S8).

### Interaction between loci explains tree size distribution in commercial pistachio orchards

In the commercial orchard dataset, two loci—one on Chromosome 3 and one on Chromosome 9—were found to be significantly associated with trunk diameter (Figure 7). We determined that this was not due to physical linkage because the markers occurred on separate chromosomal scaffolds with no evidence of misassembly at these positions (Figure 2E, Figure 4).

To dissect the interaction of these loci, we examined the most significantly associated SNP from each chromosome against each assembly. Because both SNP positions were in pseudo testcross configuration, F_1_ individuals were expected to segregate 1:1 homozygous reference: heterozygous. As such, four possible genotypic combinations existed in the F_1_ progeny, representing all possible combinations of parental chromosomes, and inheritance of these was tracked by the two combinations of F_1_ genotypes. We denote these combined genotypes {0-0, 0-1, 1-0, 1-1}, representing homozygous reference (0) or heterozygous (1) at Chr 3–Chr 9 pairs.

At each locus individually, genotype was found to be associated with trees with significantly larger (or smaller) trunk diameter, facilitating their detection in single marker association testing. Beyond this, mean phenotype between the four combinatorial genotypic classes was also found to differ significantly (Kruskal-Wallis chi-squared = 118.66, p < 0.05). Overall, the mean of genotype 1-1 was observed to be the largest in all orchards except Madera, and 1-0 the smallest, with the difference highly significant (W=29233, p<2.2e-16). In addition, we found the distribution of trunk diameter did not differ significantly when the Chromosome 9 state was 1 (i.e., 1-1 vs. 0-1), whereas it did when the state was 0 (i.e., 1-0 vs. 0-0). This result is consistent with an epistatic interaction, whereby the genotype at Chromosome 3 only affects the trait in one of the two possible Chromosome 9 backgrounds. This is evidenced by convergent/divergent rather than parallel lines when visualized using an interaction plot (Huang et al., 2012; Doust et al., 2014). We confirmed this effect with ANOVA, finding a significant interaction between the genotype at Chromosome 3 and Chromosome 9 (F= 73.573, p< 2.20E-16), although the effect would appear to be weak at Madera, which showed near-parallel lines in the interaction plot (the oldest orchard in the dataset; see Figure 8C; Table S9.1). We further repeated the same test with a Chromosome 9 marker that was both physically distant from the most significant SNP and not significantly associated in the commercial orchard association test (Pint_chr9:23111934; -log10p=1.379; ∼22 Mb from most significant SNP) and found no significant interaction between Chromosomes 3 and 9 as would be expected (F= 0.0666, p=0.7964). At all orchards, a model including an interaction term between the Chromosome 3 and 9 SNP genotypes explained a greater proportion of variance in trunk diameter than either locus individually or combined (Table S9.1).

**Figure 8.**
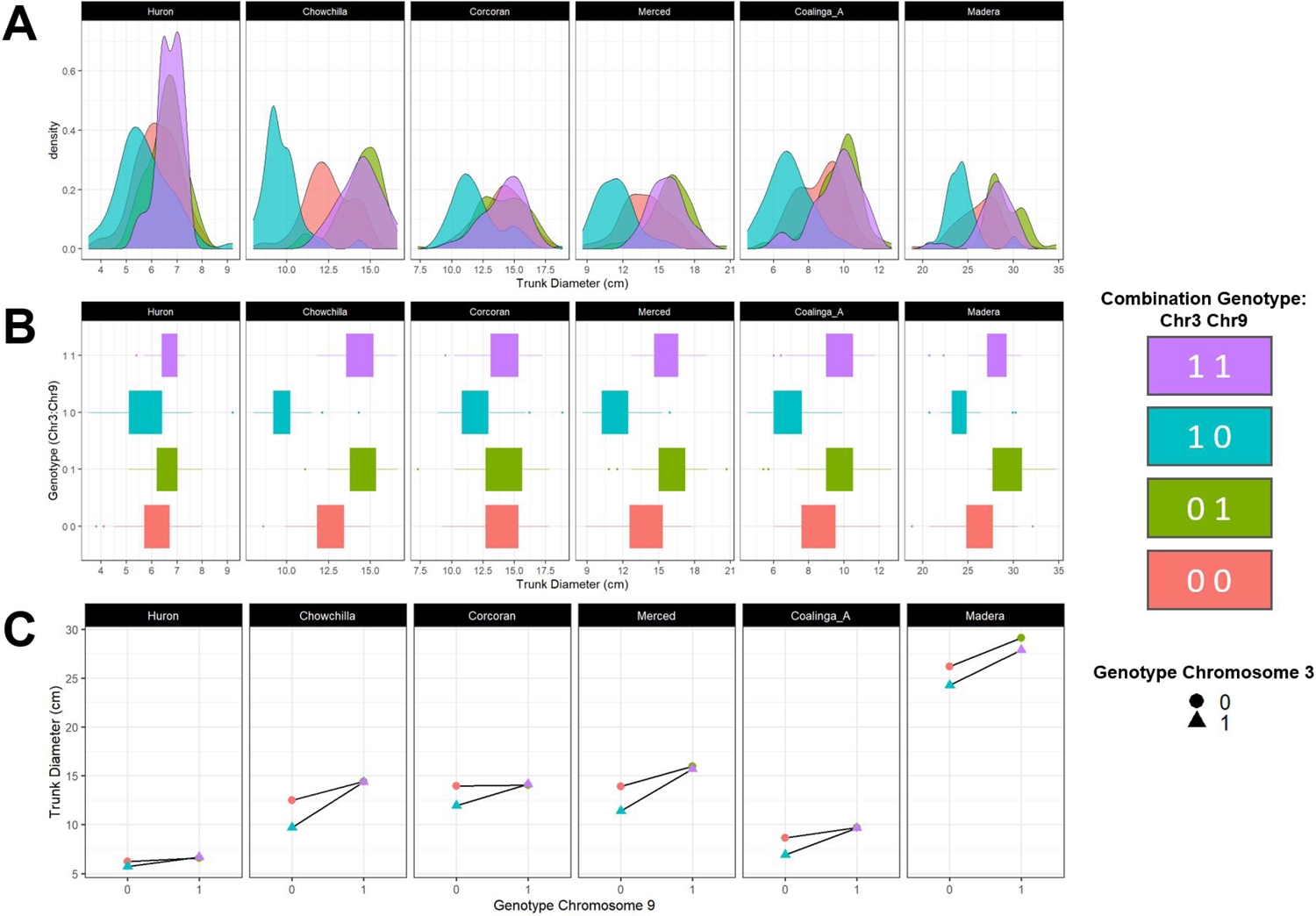
Interaction between two loci explains the size distribution of trees in commercial orchards. Panels are ordered by orchard age, youngest (left) to oldest (right). A) Density plots for trunk diameter at six commercial orchards. Each distribution is colored by one of four possible F_1_ genotypic combinations between the two loci. B) Boxplots of trunk diameter for the seven commercial orchards studied. Each box is colored by one of four possible genotypic combinations between the two loci. C) Interaction plot of trunk diameter in six commercial orchards. The x-axis represents genotype at Pint_chr9:245436, and the y-axis is trunk diameter. Triangles vs. circles represent the two alleles (1 and 0 respectively) with lines connecting the same allele at Pint_chr3:6276765 with each Chromosome 9 allele on the x-axis.

We then examined the size distribution of trees with each of the four possible genotypic combinations. Distributions of each genotypic combination were broadly found to be normal to skew-normal, although a clear bimodality was still observed in some distributions, suggesting either undetected loci or environmental factors affecting those genotypes beyond the two identified here (Figure 8A). When superimposed, these genotypic combinations described the bimodal size distribution observed in all the commercial orchards studied (Figure 8A). In all orchards except Huron, the smallest peak (mode) of trees is very well explained by a single genotypic combination (1-0) between Chromosome 3 and 9 (Figure 8A). At Huron (the youngest orchard in the dataset and where the distribution would be better described as skew-normal), this same genotypic combination simply described trees with a smaller mean trunk diameter.

### Identification of Candidate Genes

To identify candidate genes for the two loci identified in the commercial orchard dataset, we examined a 2 Mb window (± 1 Mb) surrounding the most significant SNP (-log10pmax) from each peak of markers on Chromosomes 3 and 9 and against each parental genome assembly for which the locus was identified to be segregating (see above). Each window covered a gene-rich region at each locus: there were 150 gene models within the 2 Mb window on Chromosome 3 in *P. integerrima*, and 245 genes within the Chromosome 9 window on the *P. atlantica* assembly.

Polymorphisms were detected in many of these genes; therefore, any of these genes or their promoter regions could be causal. To explore potential causal variation further, we characterized the effect of variants found in each window using SNPeff (Cingolani et al., 2012). By using genotyping of each parental species relative to each assembly, we captured additional variant positions not found in the reduced representation (GBS) sequencing of the F_1_ progeny. Because we had previously identified in which parent each trait-associated marker was segregating, we focused only on the SNPs with the expected parental segregation pattern associated with that peak of markers (*P. integerrima* at Chromosome 3 and *P. atlantica* at Chromosome 9). While acknowledging that critical variation might be overlooked, we prioritized candidate SNPs, by restricting this list to only those variants with a high predicted effect using SNPeff. Predicted high effects were diverse including start lost and stop gained, frameshift variants, and splice site modifications. We annotated genes predicted to be affected by these variants with the best hit to Arabidopsis. A full list of these candidate variants, genes, and predicted function is available in Tables S10.1 and S10.2.

Within the target window on Chromosome 3 of the *P. integerrima* assembly, we identified 114 variant positions segregating in the *P. integerrima* but not *P. atlantica* parent. These variants were predicted to have a SNPeff impact of ‘high’ on 51 gene models. Of these genes, 22 were predicted to have stop codon gaining variants and may represent particularly strong candidates. Notably, at ∼600 kb downstream from the variant most significantly associated with trunk diameter, there was a group of variants encoding premature stop codons in Pint04264. This gene showed strong sequence homology to *NB-ARC DOMAIN CONTAINING DISEASE RESISTANCE PROTEIN* (NP_194449.1), an Arabidopsis NBARC domain-containing disease resistance protein. Furthermore at only ∼45 kb downstream of the most significantly associated variant, we identified a stop-codon gaining variant in Pint04205, a gene showing homology to Arabidopsis NUCLEOPORIN 155 (NP_172938.2). In Arabidopsis, nucleoporin 155 may have a key role in effector triggered immunity (Gu et al., 2016).

Within the target window on Chromosome 9 of the *P. atlantica* assembly, we identified 117 variant positions segregating in the *P. integerrima* but not *P. atlantica* parent. These variants were predicted to have a SNPeff impact of ‘high’ on 61 gene models. Of these genes, seven were predicted to have stop codon gaining variants. Notably, at ∼100 kb downstream from the most significantly associated variant, Patl3143 contained a stop codon gaining mutation. This gene showed strong sequence homology to Arabidopsis SHORT-ROOT INTERACTING EMBRYONIC LETHAL (SIEL; NP_187492.2), a gene known to be involved in root patterning (Koizumi et al., 2011). At ∼800 kb downstream, Patl31548 contained a stop codon encoding variant and exhibited strong homology to Arabidopsis *PATTERN-TRIGGERED IMMUNITY (PTI) COMPROMISED RECEPTOR-LIKE CYTOPLASMIC KINASE 1 (PCRK1; NP_974270.1)*, a gene whose homologs are associated with both pathogen susceptibility (Sreekanta et al., 2015) and growth (Muto et al., 2011) in Arabidopsis.

## Discussion

We assembled high-quality chromosome-scale reference genomes for *P. integerrima* and *P. atlantica*, the parents of UCB-1, the most widely planted pistachio hybrid F_1_ seedling rootstock in the US. High heterozygosity and repeat content have historically rendered plant genomes difficult to assemble; we integrated whole genome short linked reads (10X Chromium), Hi-C chromosome conformation capture (Dovetail Hi-C), and whole genome long read (Oxford Nanopore Promethion) technologies to achieve highly contiguous chromosomal scaffolds exceeding an L50 of 20 Mb (Figure 2; Table S3). Quality assessment, including BUSCO and Hi-C contact, confirmed the assemblies to be of high quality and contiguous (Figure 2C, 2D, 2E; Table S4). Nucmer alignment of these two assemblies revealed the two species to be highly syntenic, with the exception of Chromosome 14 (Figure 2B). Unlike all other chromosomes, which exhibited a 1:1 relationship and conserved synteny along their entire length, *P. integerrima* Chromosome 14 exhibited strong homology to two chromosomal super-scaffolds in the *P. atlantica* assembly (denoted 14.1 and 14.2) as well as poorly conserved synteny. Subsequent construction of two high density linkage maps for each parental species (>10,000 markers) confirmed the high contiguity and accuracy of these assemblies, with the exception of Chromosome 14.

Linkage analysis revealed extremely limited recombination along the length of Chromosome 14 in both species, as well as collapsing the 14.1 and 14.2 chromosomal scaffolds to a single linkage group (Figure 3). This is consistent with the previous observation of a non-recombining pair of chromosomes that carry the sex-determining locus in the genus *Pistacia* (Kafkas et al., 2015; Sola-Campoy et al., 2015). Single marker association tests also showed sex (male or female) to be uniquely associated with Chromosome 14 (Figure 5). In the case of the *P. atlantica*, Chromosome 14 was assembled as two chromosomal superscaffolds, both containing a large number of sex-associated markers (Figure 5). These results support previous findings that sex-associated markers are heterozygous in females and homozygous in males of *P. vera* (Kafkas et al., 2015). These results strongly suggest that Chromosome 14 is the non-recombining sex chromosome pair (Sola-Campoy et al., 2015), with *P. integerimma* presenting the ZZ (male) haplotype. While our data are consistent with having recovered the Z and W haplotypes in *P. atlantica*, sex chromosomes are expected to become highly repetitive in the presence of limited recombination (Renner and Muller, 2021) and so the poor synteny (Figure 2B) may also reflect some misassembly of this chromosome. Due to the extremely limited recombination, it is challenging to define a narrow chromosomal window or candidate genes for sex determination.

We identified a simple genetic basis for vigor in grafted commercial orchards with UCB-1 seedling rootstocks. This was achieved by single marker association testing of 2,358 F_1_ UCB-1 rootstocks in an experimental and six commercial orchards. In the commercial orchard dataset, we identified two loci associated with trunk diameter (Figure 7). The same two loci were detected in measures of trunk diameter in the experimental orchard data, in addition to other minor loci (Figure 6). However, only markers on Chromosome 9 explained more than 10% of the variance in phenotype in either dataset, with the greatest observed markers explaining ∼22% of the variance in trunk diameter (Table S8, S9). The same locus on Chromosome 9 was associated with all measured architectural phenotypes in the experimental orchard except trunk diameter in the first year. Furthermore, all of the architectural traits were strongly correlated after year one (Figure 4B). Therefore, trunk diameter was taken as an indicator of overall vigor.

In the commercial orchard dataset, the proportion of variance explained by the two loci varied significantly between orchards, with the greatest predictive power observed at Chowchilla. At Corcoran, the proportion of explained variance was low for both loci; this was interesting because it was neither the oldest nor youngest orchard. Given that the same loci were detected in the ungrafted experimental orchard, the observed differences between orchards could be due to phenotypic sensitivity to either biotic or abiotic environmental factors.

In the experimental orchard dataset, associations uniquely related to year 1 traits were found on multiple chromosomes (Figure 6). A notable peak of associated markers was found on Chromosome 3, which was distinct from that detected with other traits due to its location and segregation pattern. An additional peak of markers strongly associated with trunk diameter in year 1 only was found on Chromosome 15, although the explained variance for any given marker at this locus was low. Because trees phenotyped in year 1 were measured shortly after planting, these associations may represent either a phenotype associated with the young seedling stage or greenhouse conditions where seedlings were initially propagated. Therefore, association testing of young seedlings may not be an effective strategy for improving the phenotypes of mature trees in grafted pistachio orchards.

Due to utilizing markers in testcross configuration, we were able to classify all markers in all tests by their parental segregation pattern (Figures 6, 7). In the experimental orchard data, we observed that most detected loci were segregating in the female parent, *P. atlantica*, but not *P. integerrima* (Figure 6). This is consistent with *P. atlantica* being the more heterozygous parent (1.29% vs 0.64%; Table S2). In addition, we found that the two strongly associated trunk diameter loci in the commercial orchards are segregating in opposite parents: the associated locus at Chromosome 3 segregates in the gametes of *P. integerrima* and the associated locus on Chromosome 9 segregates in the gametes of *P. atlantica* (Figure 7).

In the case of the commercial orchards, the two detected loci (on Chromosomes 3 and 9) were found to not only be highly significant, but also interacting. The genotype of the Chromosome 3 locus only influenced trunk diameter when paired with one of the two possible Chromosome 9 genotypes, a finding consistent with epistasis (Figure 8). As we do not know the parental distributions of tree size, it is unclear which genotypic combination is unexpectedly large or small. Our data are consistent with one genotype being smaller than expected or one specific combination being larger than expected. In either case, the size distribution of these four genotypic classes explains a second peak of tree size (mode) observed in commercial orchards (Figure 8A). This situation may be similar to the genetic mechanism of dwarfing rootstocks in apple and pear (Webber, 1932; Foster et al., 2015; Knable et al., 2015).

Precocity is an important trait because pistachio orchards take at least six years to become productive and nine before yield potential is realized. Analysis of when trees reached maturity in the experimental orchard revealed two loci segregating in UCB-1 together controlling ∼15% of variance for the length of juvenility (Table S7). One of these two loci was the same as the locus for vigor; bigger trees matured earlier. The other was a locus that was independent of vigor (Figure 6). Rootstocks can impact transcriptional profiles and horticultural traits including size of scions, for example in apple (Jensen et al., 2010). It remains to be determined, however, whether segregation of this second locus in UCB-1 rootstocks influences juvenility of a clonal scion or whether this locus controls precocity in *P. vera*.

Markers strongly associated with vigor at the two major loci identified gene rich regions. UCB-1 was generated to combine cold and Verticillium tolerance from different parents. The two highly significant loci segregated in the gametes of opposite parents; therefore, it is tempting to posit that these loci may encode genes associated with these phenotypes. At both loci, we found candidate genes related to the development of immune responses and stress pathways. Due to the differences in effect size between orchards, it is clear that tree size and vigor are influenced by the environment such as the presence or absence of a pathogen, edaphic factors, or treatment regime applied by the grower (such as watering or nitrogen fertilizer application). More functional work is needed to fine map these regions and validate candidates. The presented resources provide the foundation needed to accelerate such studies in pistachio.

Our findings have significant implications for pistachio growers. Phenotyping the ungrafted experimental orchard over multiple years confirmed that early phenotypic measures are poor predictors of later performance (Jacygrad et al., 2020). Therefore, roguing young trees in nurseries on the basis of phenotype does not result in more uniform orchards. In commercial orchards, a bimodal distribution of tree size has been consistently reported by growers. The identification of a simple genetic basis for this bimodality provides the opportunity to rogue seedlings using molecular markers in nurseries prior to grafting and planting in orchards. This should allow the removal of seedlings that would grow into small trees and avoid the current bimodal distributions of tree size. Marker-assisted selection of earlier maturity is also feasible. This would provide greater yields due to the larger trees as well as significant savings because smaller trees are often removed and replaced. Markers for shorter juvenility would result in orchards that would become profitable more quickly.

Tree size and vigor is often considered to be a complex trait, with interacting contributions from both genotype and environment. Consistent with this, a recent large-scale analysis of F_1_ poplar trees failed to clearly identify genetic determinants for tree size (Chhetri et al., 2020). However, rootstocks that significantly affect tree size and vigor have been identified in several horticultural woody tree systems and work has begun to explore their mechanism of action, although the specific mechanisms and genes remain elusive (Hayat et al., 2021). A previous genetic study of rootstock control of tree size in apple identified two interacting loci that resulted in dwarfing rootstocks (Foster et al., 2015; Hall et al., 2016). Similarly, a single locus (*Dw1*) was found to explain a large proportion of the phenotypic variation in apple tree size (57.2%) that also influenced flowering of the scion (Foster et al., 2015). In each case, only broad QTLs were identified. Our identification of two loci strongly associated with vigor and overall tree size in grafted pistachio is similar to the findings of Foster et al. (2015); we also found that tree size differences between genotypes became more pronounced over time, from two-years old and up. Multiple mechanisms have been proposed for this phenomenon, including altered canopy structure and leaf area, altered shoot and stem morphology, and altered hydraulic conductance (Foster et al., 2015; Hayat et al., 2021). Further work is required to identify the precise genic and physiological basis of vigor in grafted pistachio for which our findings provide the foundation.

Pistachio is an important nut crop for which genomic resources are currently limited. The UCB-1 hybrid is the main pistachio rootstock used in the US. Our results provide the genomic foundation for genetics, breeding, and marker-assisted selection of superior UCB-1 rootstock. Identifying the genetic basis and behavior of phenotypes in interspecific crosses is a central question to plant breeding and genetics and limited examples currently exist in the literature for woody plants. These resources establish the basis for future studies in this F_1_ population, of which millions of full sibling UCB-1 rootstocks are growing in the California Central Valley across multiple environments, including genotype x environment interactions and epigenetic changes in a perennial woody species. Also, because this is the first time either species has been assembled to chromosome-scale, these two assemblies provide a useful resource for broader studies in the genus *Pistacia* and the family Anacardiaceae, which includes both cashew and mango.

## Acknowledgements

We thank members of the Michelmore lab for help gathering field data, Manjula Govindarajulu and Lutz Froenicke for generating RNAseq data, Alex Kozik for data curation, Huaqin Xu for data submission to NCBI, Rick Kesseli for careful reading of the manuscript and suggestions, and Elizabeth Georgian for thorough editing. We are grateful to Bill Seaman, Ron Finucci, David Boyett, Roy Catania, Rito Zermeno, Mike Gianini, Dan Waterhouse, Carl Fanucchci, Kevin Visser, Jeff Schmiederer for their help and access to their orchards.

## Data availability

Submission to NCBI underway. Details to be provided once complete.

## Supplementary Table Legends

**Table S1.** Details of external data used in this study.

**Table S2.** Estimates of genome size and heterozygosity from genomescope and flowcytometry.

**Table S3.1** Full contig and scaffold statistics for each stage of the assembly process.

**Table S3.2.** BUSCO gene set completeness statistics for each assembly and transcriptome.

**Tables S4.1 and S4.2.** Marey maps for *P. integerrima* and *P. atlantica*, respectively.

**Table S5.** Complete phenotyping data set as used in this study.

**Tables S6.1 and S6.2.** SNPs significantly associated with sex, as detected using the *P. integerrima* and *P. atlantica* reference assemblies, respectively.

**Tables S7.1 and S7.2.** SNPs significantly associated with various phenotypes in the experimental orchard dataset, as detected using the *P. integerrima* and *P. atlantica* reference assemblies, respectively.

**Tables S8.1 and S8.2.** SNPs significantly associated with rootstock diameter in the commercial orchard dataset, as detected using the *P. integerrima* and *P. atlantica* reference assemblies, respectively.

**Table S9.1.** ANOVA results for the interaction effect of rootstock diameter associated SNPs identified on Chromosomes 3 and 9.

**Table S9.2.** η^2^ and R^2^ estimates for proportion of variance in rootstock diameter explained by SNPs identified on Chromosomes 3 and 9.

**Tables S10.1 and S10.2.** List of candidate genes and putative SNP effects for *P. integerrima* Chromosome 3 and *P. atlantica* Chromosome 9, respectively.

